# Robustness to noise reveals cross-culturally consistent properties of pitch perception for harmonic and inharmonic sounds

**DOI:** 10.64898/2026.04.15.716223

**Authors:** Malinda J. McPherson-McNato, Eduardo A. Undurraga, Aidan J. Seidle, Olivia Honeycutt, Josh H. McDermott

## Abstract

Pitch is a building block of speech and music, but the extent to which pitch perception is shared across cultures is unclear. Evidence from Western participants suggests that pitch perception relies on multiple representations. For instance, harmonic tones are easier to discriminate in noise than inharmonic tones despite comparable discrimination in quiet, suggesting that different representations are used in noise and quiet. We tested whether these effects are present cross-culturally, comparing participants from the US and a Bolivian Amazonian Indigenous community (Tsimane’). Participants heard two-note melodies and reproduced the melody by singing. Tones were either harmonic or inharmonic and were presented in noise or quiet. Both groups exhibited two characteristics of pitch perception previously seen in Western listeners: the direction of pitch changes could be reproduced with equal accuracy for harmonic and inharmonic tones in quiet but was better for harmonic than inharmonic tones in noise. However, replicating previous work, Tsimane’ vocal reproductions were much less likely to be related to the absolute pitch or chroma of the stimulus notes, differing from the tendency seen in Western participants to match pitch and/or chroma. Pitch and chroma matching behavior were more prominent in a subset of Tsimane’ whose responses to a demographic survey suggested greater integration with global and Bolivian markets and culture. The results demonstrate that the basic structure of pitch perception is shared across cultures despite other differences in pitch-related behavior that are plausibly driven by culture-specific experience.

## Introduction

Music is ubiquitous in societies around the world (Savage et al., 2015; Mehr et al., 2019). Despite profound differences in music from different places, some features are shared. One element of music that tends to be present across cultures is the use of pitch variation. Changes in pitch in a single voice or instrument convey melody, and multiple simultaneous voices or instruments can produce harmony. Pitch is also critical for conveying emotion, inflection, and (in some languages) lexical meaning in speech.

Sounds said to have pitch are typically generated by processes that repeat over time at a regular rate, such as the vibration of vocal cords, or reeds and strings on instruments. Such sounds are periodic in the time domain and have frequency spectra that are harmonic, with constituent frequencies that are multiples of a common fundamental frequency (f0; the rate of repetition of the sound). In the scientific literature, pitch has classically been defined as the percept corresponding to the f0 (de Cheveigne, 2005; Plack et al., 2005). However, recent work has suggested that pitch comparisons can be made without reference to fundamental frequency. Specifically, in some conditions, relative pitch judgments (assessing changes in pitch) are readily made with inharmonic sounds generated by perturbing the frequencies of a harmonic sound, and that lack a consistent f0 as a result (Faulkner, 1985; Moore & Glasberg, 1990; Micheyl et al., 2012; McPherson & McDermott, 2018, 2020; McPherson et al., 2022; McPherson & McDermott, 2023). This research has indicated that multiple representations underlie pitch perception depending on the circumstances: listeners either estimate and compare the f0s of sounds, or compare individual frequencies of sounds. Listeners appear to use representations of the f0 when sounds are presented in noise or when pitch must be stored over time, but instead use representations of individual frequency components when sounds are presented in quiet and with no intervening delay. Listeners evidently move fluidly and subconsciously between these two representational modes depending on the context. However, this research has only been conducted in Western participants, so the generalizability of these findings is unclear. Tests of cross-cultural consistency of pitch perception could help clarify the extent to which this aspect of perception depends on culture-specific experience, which in turn might clarify the cross-cultural consistency of pitch-related structural features of music.

In a parallel line of research, cross-cultural work comparing participants in the US to Tsimane’, a group of hunter-agriculturalists living in the Bolivian Amazon (Fig. 1a), has suggested that pitch-related behavior varies in at least some respects across populations (Jacoby et al., 2019). When asked to reproduce sequential tones by singing, US listeners tend to sing back notes that are related to the stimulus notes by integer numbers of octaves (i.e., differing by a frequency ratio that is a power of two), suggesting that notes related by octave ratios are treated as musically equivalent by US listeners (“octave equivalence”). By contrast, Tsimane’ listeners were found to exhibit no such tendency to match note “chroma”. Tsimane’ also showed less tendency to pitch match – sing back notes whose f0 matched the stimulus frequency – than US listeners. These results left open the extent to which pitch perception varies in other respects across cultures. The goal of this paper was to address this issue by assessing other aspects of pitch perception cross-culturally, comparing US to Tsimane’ listeners. We chose Tsimane’ as a comparison group given that they are one culture for whom there is some evidence for pitch-related behavior that differs from that seen in Western listeners.

**Figure 1.**
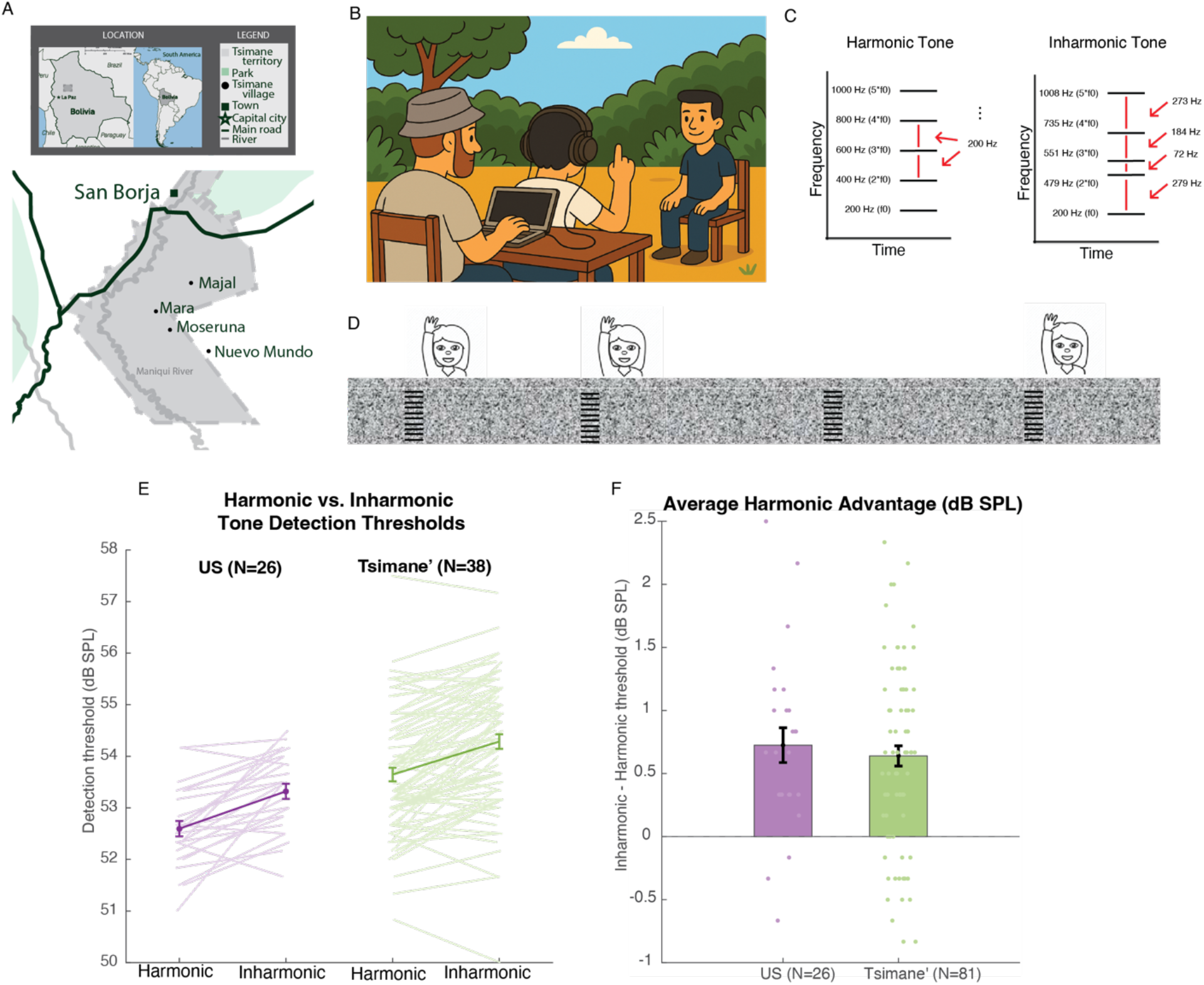
Harmonic advantage for detecting sounds in noise is present cross-culturally. A. Map of the region in Bolivia where testing took place, showing the four villages where data were collected (Majal, Mara, Moseruna and Nuevo Mundo). B. Example experiment setup for Experiment 1. Participants were facing away from the experimenter (left), and raised their hand when they heard a tone. For all experiments in Bolivia, translators (right) explained experiment instructions in Tsimane’. C. Comparison of sample Harmonic and Inharmonic stimuli. Inharmonic stimuli were generated by randomly jittering the frequency above the fundamental (f0) by up to 50% of the f0. D. Schematic of Experiment 1 setup. Participants heard continuous noise and were asked to raise their hand every time they heard a tone. The levels of the tones were varied by the experimenter to find the lowest detectable level for each participant. E. Results for Experiment 1 showing the individual and group average detection threshold. F. Results for Experiment 1 showing that both groups are better at detecting harmonic vs. inharmonic tones in noise.

To probe representations of pitch, we developed cross-cultural versions of experiments previously run on US participants. We first asked participants to detect harmonic and inharmonic tones in noise by raising their hand when they heard a tone. In a second experiment, we asked participants to sing back short melodies composed of either harmonic or inharmonic tones, played in background noise at two different signal-to-noise ratios (SNRs). The SNRs were chosen so that at the Low SNR, US participants would exhibit harmonic advantages in discrimination, but at the High SNR, they would have comparable discrimination between harmonic and inharmonic stimuli. Our primary question was whether Tsimane’ listeners would exhibit comparable sensitivity to harmonic structure, as measured by better detection and discrimination of harmonic than inharmonic tones in noise, despite failing to exhibit pitch and chroma matching. We also asked whether the direction accuracy of sung responses would be comparable for harmonic and inharmonic tones without noise, as would be expected to occur for US listeners.

A priori, it was not clear what to expect from the harmonic/inharmonic manipulations. Some reason to expect a harmonic advantage like that seen in Westerners comes from previous evidence that Tsimane’ are sensitive to harmonic structure when asked to perceptually segregate multiple concurrent tones (McDermott et al., 2016; McPherson et al., 2020). On the other hand, to pitch- or chroma-match, it is almost certainly necessary for listeners to maintain a representation of the absolute pitch, i.e. of the f0 (Van Hedger et al., 2015). The absence of pitch and chroma matching is thus consistent with the absence of representations of the f0 (though it could also be explained by listeners not having learned to coordinate singing with heard pitches).

## Results

### Experiment 1: Advantage for detecting harmonic tones in noise is present cross-culturally

The purpose of Experiment 1 was to test whether Tsimane’ participants exhibit one known signature of sensitivity to harmonic structure: the advantage for detecting harmonic sounds in noise, previously seen in US participants (McPherson et al., 2022). Additionally, Experiment 1 served as a control to ensure that all stimuli presented in Experiment 2 were audible. Participants heard continuous Threshold Equalizing Noise presented at 65 dB SPL, and were instructed to raise their hand every time they heard a tone (Fig. 1b-c). Participants faced away from the experimenter to avoid visual cues to the tone presentation timing. The experimenter adjusted the intensity of the tones to determine the faintest tone the participant could reliably detect. The experimenter was blind to whether the presented tones were Harmonic or Inharmonic.

Despite overall differences in average thresholds, we found that both Tsimane’ and US participants were sensitive to harmonic structure: detection in noise was better for harmonic compared to inharmonic tones for most participants in each group (Fig. 1d). There was a significant effect of harmonicity at the group level (F(1,105)=71.57, p<.0001, ηp²=.41), and no interaction between the effect of harmonicity and the participant group (F(1,105)=0.27, p=.60, ηp²=.003, BF10=0.26, providing moderate evidence for the null hypothesis). These results suggest that listeners in both cultures are sensitive to harmonicity, and can use harmonic structure to help segregate sounds in noise, potentially using the pitch cue present in a representation of the f0 to aid detection.

### Experiment 2: Harmonic advantage for discrimination in noise is present cross-culturally

Participants were asked to sing back two-note melodies presented in noise (Fig. 2a-b). We recorded their sung responses, extracted the f0s of the two reproduced notes, calculated the direction of pitch change between reproduced notes, and compared the direction of pitch change (up or down) to that of the stimulus (as an indirect measure of pitch perception). The sound levels of the tones were varied to create either a High SNR, for which the notes were easy to hear, or a Low SNR, for which the notes were harder to hear. The experiment was split into four blocks. Two blocks contained Harmonic tones and the other two contained Inharmonic tones, crossed with High and Low SNRs. The noise was played at a continuous level of 65 dB SPL. The tones were presented at 57 dB SPL (-8 dB SNR) and 70 dB SPL (+5 dB SNR) in the Low and High SNR conditions, respectively. We ran a supplemental experiment in US participants (Supplemental Experiment 1, Supplemental Fig. 1) to confirm that these SNRs would produce the expected Harmonic advantage in the Low SNR condition, but not in the high SNR condition. Moreover, the detection results in Experiment 1 indicate that the tones in the Low SNR conditions were presented, on average, 3-4 dB above their detection thresholds, making them readily audible in all cases. The main question was whether Tsimane’ would also exhibit this effect.

**Figure 2:**
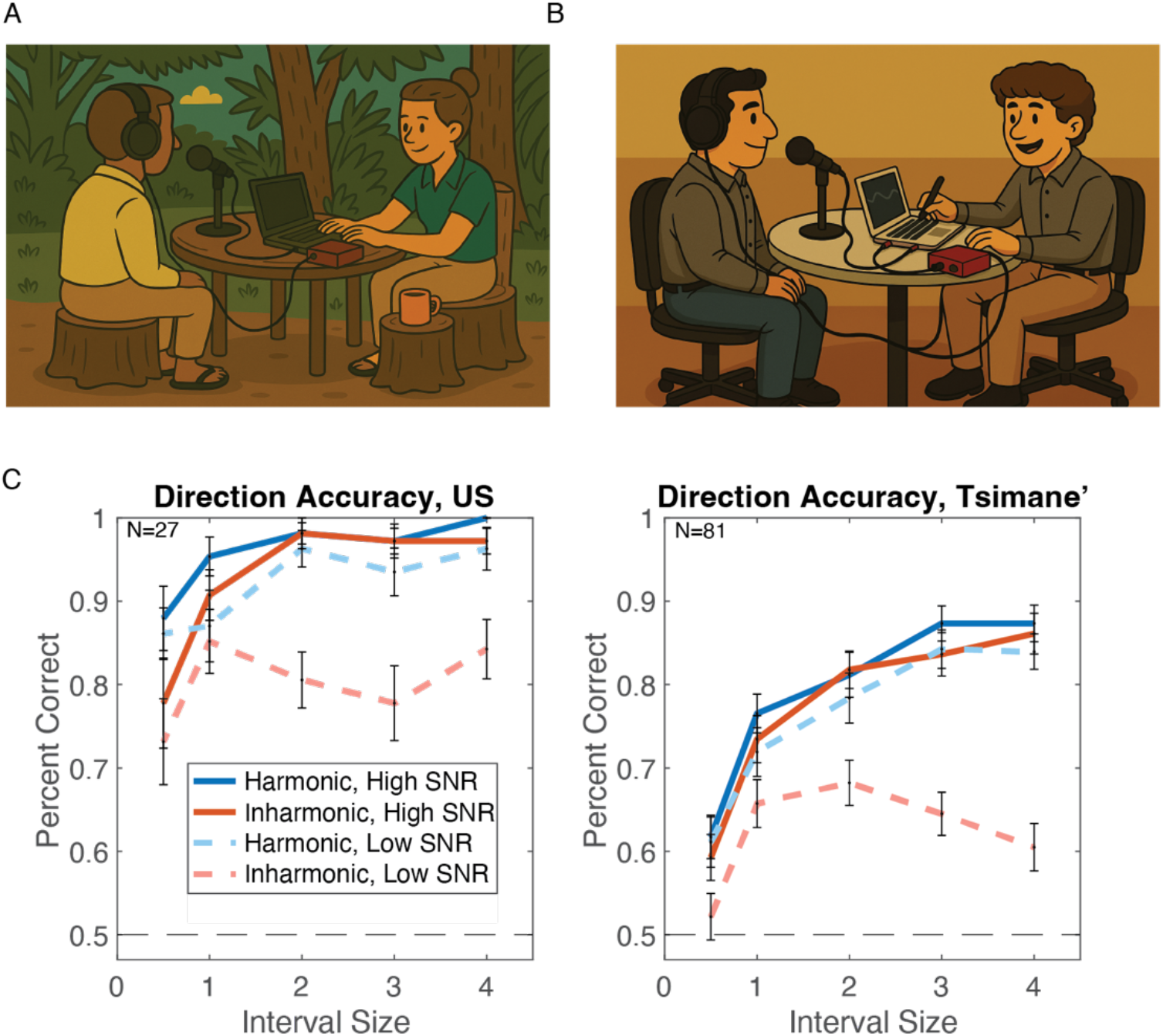
Performance is worse for Inharmonic tones at Low SNRs for both participant groups. A. Experiment setup for Experiment 2 in Bolivia. Participants sit facing the experimenter, wearing headphones. A microphone was positioned in a comfortable place in front of the participant. The experimenter played sounds using a MacBook Air laptop. B. Experiment setup for Experiment 2 in the US. Experiments were conducted in quiet, but not soundproof, rooms. C. Direction accuracy as a function of interval size for US (left) and Tsimane’ (right) participants. Results were averaged across positive and negative intervals (e.g. direction accuracy for -1 and 1 are averaged in the current plots). Error bars show SEM.

In each block, tone fundamental frequencies were either in the participant’s singing range (defined based on their sex) or outside their singing range (below the singing range for female participants, or above for male). The order of the blocks was randomized, and experimenters were blind to which block was being presented.

#### Cross-cultural similarities in reliance on harmonic structure in noise

Despite a large effect of Group on direction accuracy (F(1,106)=34.9, p<.0001, ηp²=.25, BF10= 3.97 × 10⁴), with US more accurate than Tsimane’, the effect of the stimulus condition on accuracy was quite similar across groups. At the High SNR, sung direction accuracy was comparable for Harmonic and Inharmonic tones in both groups (F(1,106)=3.87, p=.052, ηp²=.04, BF10=0.091), showing moderately strong evidence for the null hypothesis (Rouder et al., 2009), Fig. 3c, and no significant interaction between Group and Harmonicity: (F(1,106)=0.31, p=.58, ηp²=.003, BF10=0.16), indicating moderate evidence for the null hypothesis. There was a modest effect of harmonicity in the US group (F(1,26)=6.73, p=.02, ηp²=.21); this was driven by the lowest frequency difference (no effect of harmonicity with only four higher steps, F(1,26)=3.14, p=.09, ηp²=.11, BF10=0.69, showing anecdotal evidence for the null), and may be accounted for by pitch and chroma matching that occur in US participants when singing (see below). The US participants were close to ceiling for larger step sizes, but overall the similarity in performance for Harmonic and Inharmonic tones is consistent with the supplementary experiment used to select the SNRs (Supplemental Fig. 1b), and with extensive prior evidence in US listeners that discrimination in quiet is indistinguishable for harmonic and inharmonic tones (Faulkner, 1985; Micheyl et al., 2012; McPherson & McDermott, 2018, 2020; McPherson et al., 2022; McPherson & McDermott, 2023). Thus, the critical result for these conditions is the similar performance for Harmonic and Inharmonic conditions in Tsimane’ (F(1,80)=1.64, p=.20, ηp²=.02, BF10=0.23), showing moderate support for the null), for whom traditional psychoacoustic tasks are less viable.

**Figure 3:**
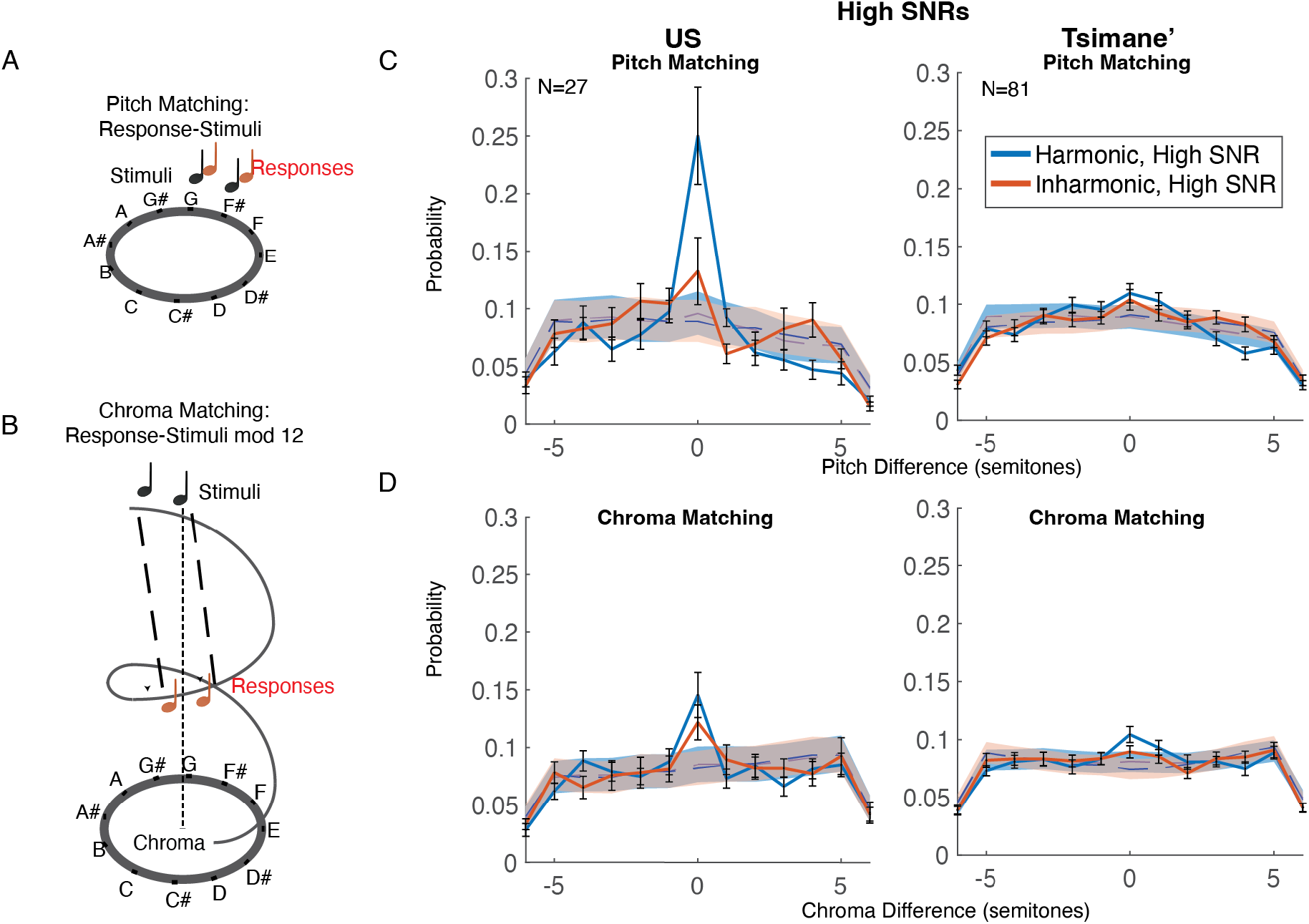
Pitch and Chroma Matching are stronger in US than in Tsimane’. A. Schematic of pitch matching analysis. B. Schematic of chroma matching analysis. To determine the ‘chroma difference’, we subtracted the response from the stimulus after transposing the response into the singing range, yielding a difference between the stimulus and response modulo 12. C. Histograms of stimulus-response pitch differences for High SNR conditions, separated by participant group. Error bars plot SEM across participants, calculated via bootstrap. The shaded regions denote the null distributions, computed from histogram of permuted datasets where stimulus-response correspondence was randomized. D. Same as C, but for stimulus-response chroma differences.

A different pattern of results was evident at the Low SNR. Here, both groups showed much better direction accuracy for Harmonic than Inharmonic tones (Main effect of Harmonicity: F(1,106)=54.10, p<.0001, ηp²=.34). The effect was similar for the two groups, with no significant interaction between Group and Harmonicity: F(1,106)=0.35, p=.55, ηp²=.003, BF10=0.18, showing moderate evidence for the null hypothesis. This pattern of results was further substantiated by an analysis of the mean produced interval as a function of the stimulus interval, which for both groups showed less accurate reproductions specifically for inharmonic tones at the low SNR (Supplemental Fig. 2).

These results suggest that, like US listeners, Tsimane’ use at least two different representations when making pitch judgments, and that they use them in roughly the same way as US listeners. In particular, they show signs of relying on individual frequency components at High SNRs, and on a representation of the f0 at Low SNRs.

#### Cross-cultural differences in pitch- and chroma-matching

Using the same sung responses from Experiment 2, we analyzed pitch and chroma (octave) matching for all conditions (3a-b, and Supplemental Fig. 3). Here, instead of comparing the f0 change between sung notes to the f0 change of the stimulus, we calculated the difference between the f0s of the stimulus and response in semitones, expressed this difference modulo 12. We then split responses by register. If the listener sang within 6 semitones of the target, their responses were included in the pitch matching analysis. Trials in which the response fell more than 6 semitones away from the stimulus were instead included in the chroma matching analysis. We summarized the results in histograms (see Methods for details of how histograms and null distributions were obtained). For Inharmonic tones (which do not have a coherent f0), we used the lowest frequency component in lieu of the f0, as it is in principle possible to hear out this frequency component and match a sung note to it (McPherson & McDermott, 2018). Pitch and chroma matching are evident as peaks at 0, indicating that the participant matched the pitch or chroma of the stimulus notes. Here we present results for the High SNR conditions, as this is most similar to the previous experiment that was analyzed in this way (Jacoby et al., 2019). Results were similar for the Low SNR (Supplemental Fig. 3).

As expected, US-based participants showed significant pitch and chroma matching for harmonic tones (Pitch, test for significance above the null distribution in the histogram bin that indicates pitch/chroma matching around 0 semitones: p<.0001; Chroma: p<.0001; Fig. 4c-d). They also showed a significant tendency to pitch- and chroma-match to the lowest frequency component of the inharmonic tones (p<.0001 in both cases), but this was significantly weaker than for harmonic tones (main effect of harmonic vs. inharmonic, F(1,26)=9.45, p=.0049, ηp²=.27). This difference is expected given that inharmonic tones yield weaker representations of absolute pitch (McPherson & McDermott, 2018). We note that the harmonic-inharmonic difference in pitch matching likely explains the slight harmonic advantage seen in sung direction accuracy for US participants at high SNRs in Fig. 3c. Pitch matching would indirectly serve to increase the direction accuracy of the sung responses, increasing accuracy for harmonic compared to inharmonic stimuli.

**Figure 4:**
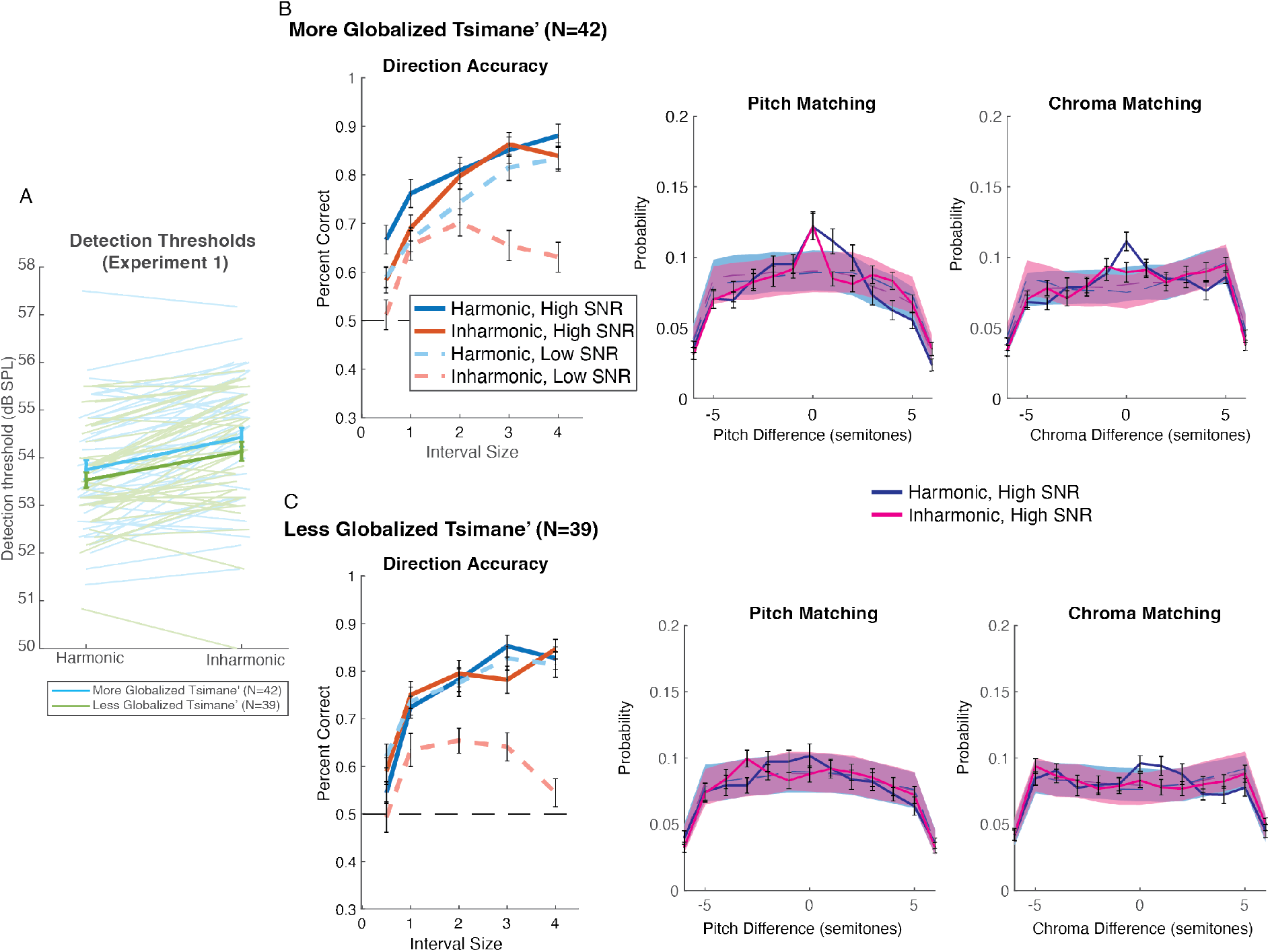
Effects of global integration on pitch- and chroma-matching. We quantified exposure to globalized markets and culture using an index that aggregated a set of demographic variables (see Methods). We divided the Tsimane’ participants into two groups using a median split and then separately analyzed the data from Experiments 1 and 2. (A) There was no significant difference in detection thresholds for More and Less Globalized Tsimane’. Error bars show standard error of the mean. More Globalized Tsimane’ (B) showed much greater tendency to pitch and chroma match than Less Globalized Tsimane’ (C). By contrast, both groups showed similar direction accuracy for Harmonic and Inharmonic tones at the High SNR, but a pronounced harmonic advantage at the Low SNR. These results provide a further dissociation between pitch phenomena, while also providing some evidence that increases in pitch and chroma matching in Tsimane’ are resulting from changes in the extent to which they are exposed to global culture.

As in a previous study, Tsimane’ participants showed much less tendency to match the pitch or chroma of harmonic tones, though they did do so to a degree that was above chance levels (Pitch: p=.0001; Chroma: p<.0001). This effect was somewhat weaker for Inharmonic tones, though this difference was not statistically significant (p=.11). To compare across groups, we analyzed the value of the histogram bin around 0 semitones for both pitch and chroma. There was a significant main effect of Group for Pitch (F(1,106)=12.95, p=.0005, ηp²=.11) and Chroma (F(1,106)=9.20, p=.003, ηp²=.08), confirming that pitch and chroma matching are less pronounced in Tsimane’ than in US non-musicians.

#### Cross-cultural differences are not explained by differences in sensitivity

Could any of the effects in Experiment 2 (Figures 2 or 3) be explained by the overall difference in thresholds seen in Experiment 1 (Figure 1)? The main consequence of differences in overall thresholds is that the tones of Experiment 2 could have been closer to threshold for Tsimane’ participants, on average, despite being presented at the same physical SNR. A priori it seemed unlikely that such a difference could explain the effects of interest in Experiment 2. First, the effects related to direction accuracy involve similarities between groups. Second, the effects related to pitch and chroma matching were substantiated using the high SNR condition, in which the tones were far above threshold for both groups (presented at 70 dB SPL, with thresholds in the range of 52-54 dB SPL, depending on the group and harmonicity condition).

We nonetheless addressed the issue directly, reanalyzing the data from Experiment 2 after subsampling participants to equate overall detection thresholds in Experiment 1. We split the data in Experiment 1 for each participant, using one half of the data to select a subset of Tsimane’ participants whose mean thresholds were close to the mean thresholds of the US participants. We then used the second half of the data to verify that the thresholds were indeed approximately matched (53.3 dB SPL for the Tsimane’ subset, N=38, compared to 53.4 dB SPL for the US group, N=26; not significantly different, t(62)=0.15, p=.88, BF10=0.26, showing moderate evidence for the null). When we analyzed the results of Experiment 2 for just this subset of participants, we found results that were similar to those of the main analysis (Supplemental Fig. 4). Specifically, using the same Group × Harmonicity × Noise mixed-model ANOVA as the main analysis, direction accuracy showed the same significant effects of harmonicity (F(1,62)=34.47, p<.0001, ηp²=.36), noise (F(1,62)=26.35, p<.0001, ηp²=.30), and their interaction (F(1,62)=15.74, p=.0002, ηp²=.20) seen in the main analysis, with no Group interactions (all p>.50) — indicating these stimulus-condition effects were consistent in US and Tsimane’ participants. The large group differences in direction accuracy (F(1,62)=21.92, p<.0001, ηp²=.26), chroma matching (F(1,62)=7.02, p=.010, ηp²=.10), and pitch matching (F(1,62)=9.41, p=.003, ηp²=.13) also remained present. Overall, these analyses suggest that a subset of the Tsimane’ participants have tone-in-noise thresholds that are on par with US participants, and that the modest differences in average thresholds do not affect the conclusions from Experiment 2.

#### Effects of global integration on pitch-related behavior

We hypothesized that the increased pitch and chroma matching seen here relative to a previous study (which failed to detect any significant effects in Tsimane’) is driven by the increased contact that many Tsimane’ have with globalized culture compared to previous years. One prediction of this hypothesis is that there should be reliable individual differences in the extent of pitch and chroma matching in individual participants. We first tested whether this was the case by correlating the extent of pitch and chroma matching for individual participants on odd versus even trials. For both groups, these scores were more reliable than would be expected by chance, demonstrating meaningful variation at the level of individual participants. Reliability was high for both measures in the US (chroma: r = .93, 95% CI [.86, .97]; pitch: r = .92, 95% CI [.82, .96]). Reliability was lower in the Tsimane’ (unsurprisingly given that the overall extent of pitch and chroma matching was low) but still higher than chance (chroma: r = .35, 95% CI [.15, .53]; pitch: r = .40, 95% CI [.20, .57]).

To explore whether variation in the pitch and chroma matching of Tsimane’ participants might relate to their exposure to globalized culture, we used a survey of ethnographic variables to split participants into two groups that were more vs. less integrated with the global economy. This approach was developed (and pre-registered) in previous work which found that consonance preferences were evident in Tsimane’ with greater global economic integration (McPherson-McNato et al., 2025). Here we completed a conceptual extension of this previous experiment, hypothesizing that the more integrated group might show stronger signatures of chroma and pitch matching.

The two groups of Tsimane’ showed similar effects of harmonicity on detection and relative pitch. In Experiment 1, there was no interaction between the effect of harmonicity and the two Tsimane’ participant groups (F(1,79)=1.30, p=.26, ηp²=.02), and overall thresholds were similar (Figure 4a). In Experiment 2, both Less and More Globalized Tsimane’ showed much better direction accuracy for Harmonic than Inharmonic tones in the Low SNR condition (Main effect of Harmonicity: F(1,79)=74.52, p<.0001, ηp²=.49; Fig. 4b). The effect was similar for the two groups, with no significant interaction between Group and Harmonicity: F(1,79)=0.52, p=.47, ηp²=.007, BF10=0.18, showing moderate evidence for the null hypothesis. There was also an interaction between SNR and Harmonicity in both groups (Less Integrated: F(1,38)=24.86, p<.0001, ηp²=.40, More Integrated: F(1,41)=6.19, p=.02, ηp²=.13). These findings are consistent with the idea that any pitch representations that aid detection in noise are largely independent of global integration, and that the use of two different pitch representations is widespread.

We analyzed pitch and chroma matching in the two groups of Tsimane’ while pooling across the High and Low SNR conditions, to maximize power. This analysis revealed clear differences between groups. Less Globalized Tsimane’ participants showed a tendency to match the pitch of Harmonic tones that reached statistical significance (test for significance above the null distribution in the histogram bin centered around 0 semitones: p=.04; Fig. 4), but Pitch matching for Inharmonic tones (p=.46), Chroma matching for Harmonic tones (p=.052), and Chroma matching for Inharmonic tones all did not reach significance (p=.44). By contrast, the More Globalized Tsimane’ group exhibited significant Pitch and Chroma matching for both tone types (p=.0012 for Harmonic and Inharmonic tones for Pitch matching, p<.0001 for Harmonic tones for Chroma matching and p=.008 for Inharmonic tones for Chroma matching). The difference between the two Tsimane’ groups was reflected in a significant main effect of Group on Pitch and Chroma matching (F(1,79)=6.57, p=.01, ηp²=.08, when pooled across both SNR conditions). As with the US group, the More Globalized Tsimane’ demonstrated a modest benefit of harmonicity on direction accuracy (F(1,41)=9.58, p=.004, ηp²=.19), and this may similarly be accounted for by the modest pitch and chroma matching that occurs in these participants.

To supplement the median split analysis, we completed a second analysis using the Globalization index as a continuous regressor. Globalization significantly predicted the extent of matching for both pitch (β =0.0059, SE=.0024, t(79)=2.46, p=.017, R^2^=0.07) and chroma (β =0.0037, SE=0.0016, t(79)=2.31, p=.023, R^2^=0.06). By contrast, Globalization did not predict singing direction accuracy (R^2^=0.001, F(1,79)=0.11, β =0.0026, SE=0.0078, p=.74). The Globalization effect on pitch- and chroma-matching remained after controlling for participant age (pitch: p=.050, chroma: p=.038), which itself was not a significant predictor in either case; (p>.19). To test whether the Globalization effect was driven by a subset of highly globalized or non-integrated participants, we added a quadratic term to each regression. The quadratic term was not significant in either case (pitch: p=.34; chroma: p=.16), indicating no evidence of a nonlinear or threshold-like relationship between Global Integration and pitch- and chroma-matching. These analyses confirm the findings from the median split.

Overall, these results provide additional support for a dissociation between the basic components of pitch perception, which seem to be present across cultures and not qualitatively affected by integration with global culture, and pitch-related singing behavior, which varies substantially across cultures. The results also support the conclusion that recent globalization is having measurable effects on music-related behaviors, in that pitch and chroma matching are becoming more prevalent in individuals with higher levels of global integration.

## Discussion

We used singing to probe pitch representations cross-culturally and found evidence that the ability to use multiple pitch cues and the way in which these cues are used are shared across cultures. Moreover, these shared abilities dissociate from pitch and chroma matching, the incidence of which varied across the groups we tested.

### Multiple representations of pitch are present across cultures

Both Tsimane’ and US participants showed comparable sung direction accuracy for harmonic and inharmonic tones when the tones were presented in High SNRs. This result suggests that both groups can make relative pitch judgments without a consistent f0, for instance by registering shifts that occur between individual frequency components (McPherson & McDermott, 2018). By contrast, when tones were presented at Low SNRs, both groups showed an advantage for Harmonic over Inharmonic stimuli, suggesting that in these conditions they relied on an f0-based representation of pitch. Together, these results suggest that multiple representations of pitch are a universal feature of pitch processing.

In principle, the harmonic advantage at low SNRs might be viewed as a grouping effect, whereby harmonic tones are more readily separated from noise. However, the SNRs were chosen so that even the inharmonic tones were substantially above their detection threshold in the low SNR condition. Thus, the harmonic advantage is unlikely to be driven by detection-related effects, and instead is likely driven by the ability to estimate the direction of a pitch change from a set of frequencies. The advantage plausibly reflects the ambiguity of individual frequency components when heard in noise, which complicates the estimation of pitch direction from individual frequencies. Representations of f0 integrate across frequencies and appear to be more noise-robust as a result.

While sung reproduction is not a direct proxy for perceptual judgments of pitch, as it also involves articulation and production constraints(Dalla Bella et al., 2007; Pfordresher & Brown, 2007; Hutchins & Peretz, 2012), the direction accuracy effects were consistent across perceptual and sung tasks in US listeners. This consistency suggests that the variation in sung direction accuracy with stimulus conditions reflects perceptual representations rather than production constraints.

The cross-cultural consistency of relative pitch representations motivates analogous experiments in non-human animals to assess whether such representations are unique to humans (Walker et al., 2019; So et al., 2020). The results also motivate future experiments on the pitch representations of more extended sequences of sounds, to assess whether representations of melodic and/or prosodic contour are consistent across cultures.

### Dissociation between f0-related pitch abilities

Despite appearing to use f0-based pitch to detect sounds in noise (Experiment 1) and to extract relative pitch in noise (Experiment 2), Tsimane’ listeners do not use f0-based representations in all the settings in which US listeners do. This is evident in pitch and chroma matching behavior during singing, where Tsimane’ do not match absolute pitch when reproducing melodies. This result demonstrates that pitch and chroma matching behavior are not an inevitable consequence of Western-like pitch perception, presumably depending on some type of musical experience. One possibility is that pitch and chroma matching behavior is a habit that develops from experience singing in groups. This practice has not been widespread in traditional Tsimane’ culture (though it is changing with the spread of Christian churches in the region), whereas US residents tend to engage in group singing from an early age. Consistent with this idea, we found that the very weak tendency to pitch and chroma match was greater in Tsimane’ with higher scores on a measure of global integration (Supplementary Table 1 and Fig. 3).

The present results clarify the previously observed cross-cultural difference in pitch and chroma matching by showing that it likely does not reflect the absence of f0-based representations of pitch (Pressnitzer & Demany, 2019). However, the dependence of the effects on singing leaves open whether the differences in pitch and chroma matching reflect cross-cultural variation in perceptual octave equivalence. It remains possible that the effects are specific to singing and reflect differences in the use of pitch rather than differences in pitch itself.

### Use of singing in cross-cultural research

The present work further underscores the utility of active paradigms, such as those involving singing and tapping, in cross-cultural research (Jacoby & McDermott, 2017; Jacoby et al., 2019; Jacoby et al., 2024). The results here used an active paradigm to replicate results obtained in US listeners using more traditional psychophysical methods (McPherson et al., 2022). Active paradigms have the advantage of utilizing behaviors that appear to be universal, independent of their cultural background (Mehr et al., 2019), and are thus applicable in populations where traditional psychophysical instructions might be difficult to translate. Active paradigms also allow for the rapid collection of multi-dimensional data (here, for example, the simultaneous collection of chroma/pitch and interval direction and magnitude accuracy). These types of paradigms may also be useful for testing individuals with shorter attention spans, such as children. Such paradigms are nonetheless more complicated to interpret than traditional psychophysical methods, as the measured behavior combines effects of perception and motor production.

### Effects of global integration

We replicated an earlier study in finding significant differences between US and Tsimane’ in the tendency to pitch and chroma match, but nonetheless observed weak effects in Tsimane’ where a previous study did not (Jacoby et al., 2019). This result plausibly reflects the changes that have occurred in the area since null effects of chroma were first observed in Tsimane’ in 2017 and 2018 (Jacoby et al., 2019). Consistent with this hypothesis, we found modest differences between subsets of Tsimane’ participants when the large group was split based on a measure of global integration. This result complements a recent analysis that found analogous effects on consonance preferences in different groups of Tsimane’ (McPherson-McNato et al., 2026). These results together suggest that the globalization rapidly reshaping life experiences in small-scale societies has measurable effects on experimental results.

## Methods

### 1. Study Participant Details

We used a brief, independent “filter” test, described in #4 under *Procedure* below, to determine which participants to include in our final analyses. The filter test required participants to sing back 8 melodic intervals composed of pure tone notes, in two different pitch ranges, from which we measured sung direction accuracy. Data from participants who performed at or better than 75% accurate (12/16) on this filter test were included in further analysis. The filter task and inclusion criterion were the same for both participant groups.

Participants were also screened with a test that measured hearing thresholds for diotically presented pure tones at 10 frequencies (60, 127, 285, 605, 1358, 2155, 3229, 5126, 8137, and 11500 Hz, Supplemental Fig. 5). Participants with average thresholds (averaged over the frequencies from 60 to 8137 Hz, which covers the full range of experimental stimuli) above 50 dB SPL would have been excluded from the analysis; however, all tested participants passed this inclusion criterion.

US: 33 participants from Boston and West Lafayette, Indiana, were enrolled in Experiments 1 and 2.27 of the 33 participants (Female=15, mean age=25.5, S.D.=8.1) passed the filter test and were included in further analysis. All of these participants passed the hearing screening. One participant’s detection thresholds (Experiment 1) were incorrectly measured due to experimenter error and their data was excluded from the Experiment 1 analysis.

Tsimane’: 111 individuals were enrolled in the experiment. Three were removed because their thresholds in Experiment 1 were more than two standard deviations above (worse than) the group average. 16 did not complete Experiment 2 because they could not understand the instructions or could not complete the task (for example, did not sing loudly enough for the microphone to reliably capture their responses). One additional participant was removed because of a data write error in the field. Of the 91 remaining participants (Female=34, mean age = 28.5, S.D.=10.9), 81 passed the filter test and were included in further analysis. One of these participants did not complete the hearing screening, but all others did, and passed it. One of the main analyses split Tsimane’ participants into two groups based on demographic characteristics; the details of these groups are given below in the section *Demographic Survey in Tsimane’*.

#### Sample Sizes

For Experiment 1, we based our power estimates on previously published experiments measuring detection in noise (McPherson et al., 2022). We calculated that we needed 18 participants per group to be powered to see an effect one quarter the size of the previously observed harmonic/inharmonic distinction (ηp²=.37), at a .05 significance, 95% of the time, thus Experiment 1 was adequately powered, and the final sample size was determined by the larger sample requirements of Experiment 2.

We determined recruitment targets for Experiment 2 (and thus Experiment 1, which used the same participants) based on the sample sizes required to achieve adequate statistical power. Previously published power analyses for a similar experiment indicated that 17 participants were required to detect chroma matching on par with that seen in US participants 90% of the time (Jacoby et al., 2019). In Bolivia, we recruited a sample size substantially larger than this number in part to complete the globalization analyses in Experiment 2. In practice, in Bolivia we recruited all adults who were willing to participate in each village we visited. For US data, our final sample size exceeds the minimum required to detect a within-between group interaction with an effect size of ηp²=.05 at α=.05 with 95% power, ensuring that the study remained well-powered despite differences in group sizes.

#### Ethics

All procedures with adults were completed with approval from the Committee on the Use of Humans as Experimental Subjects at MIT. Experiments in Bolivia were also approved by the Tsimane’ Council (the governing body of Tsimane’ in the Maniqui basin, where the experiments took place). Experiments with US adults and online experiments were also approved by Purdue University’s Human Research Protection Program. Experiments were conducted with the informed consent of the participants. Participants were compensated with money (US) and packages of goods (Tsimane’).

## 2. Method Details

### Stimuli

#### Complex Tones

Experiments used complex tones containing 10 equal amplitude harmonics, added in sine phase. Tones were windowed with 10ms half-Hanning windows and were 500ms in duration. To make tones inharmonic, each frequency component (other than the f0) was ‘jittered’ by up to 50% of the f0 value. To select jitters, we sampled values from the distribution U(-.5, .5), multiplied that by the f0, then added the resulting value to the frequency of the respective frequency component. We iteratively selected jitter values, moving up the harmonic series, and rejecting jitter values that resulted in adjacent harmonics being within 30 Hz to avoid salient beating. This procedure was identical to jittering found in previous experiments (McPherson & McDermott, 2018; Popham et al., 2018; McPherson & McDermott, 2020; McPherson et al., 2022). Further characterization of these jitters, along with summed-autocorrelation analyses can be found in McPherson & McDermott (2018) and Supplemental Fig. 6. Using this jittering procedure, we randomly generated 100,000 possible jitter patterns and selected the 20 that minimized peaks in the autocorrelation function. One of these 20 possible jitter patterns was chosen per experiment per participant so that all the inharmonic trials across the experiment had the same jitter values.

The initial training sequence that introduced participants to the singing task (numbers 2-4 in *Procedure* below) for Experiment 2 used pure tones that only contained harmonic 1 of the complex tones described above, but all other parameters (windowing, duration) were identical.

#### Noise

All background noise across experiments was Threshold Equalizing (TE) Noise (Moore et al., 2000) with a minimum frequency of 20 Hz and a maximum frequency of 20,000 Hz. For Experiments 1 and 2, a 20-minute sample of TE noise was pre-generated and loaded for each experiment, and presented at 65 dB SPL. For Supplemental Experiment 1 (conducted online), each tone was embedded in a randomly generated sample of TE noise that began and ended 200ms after the tone and was windowed with a 10ms Half Hanning window. Because this experiment was conducted online, we could not control the absolute stimulus levels, but the SNRs were set in the same way as in the in-person experiments (see below).

### Sound presentation

Sounds were presented over Sennheiser HD 280 Pro circumaural headphones and sampled at 44.1 kHz. For Experiment 1, headphones were connected directly to a Mac laptop computer, and in Experiment 2 to a Focusrite Scarlett 2i2 USB sound card. Both systems had been calibrated ahead of time using a GRAS 43AG Ear & Cheek Simulator and a Svantek SVAN 977 audiometer to ensure that the tones and noise were presented at the desired levels regardless of the presentation method. Sennheiser HD 280 Pro circumaural headphones are designed to attenuate ambient noise (by up to 32 dB depending on the frequency). This attenuation, plus the continuous 65 dB SPL background noise in all the experiments, was sufficient to mask ambient background noise at the modest levels encountered in the rural testing environment. We optimized listening conditions by choosing test locations that were distant from potential acoustic disturbances like community activities, and we typically assigned a team member to keep children and animals out of earshot of the experimental stations. The attenuation of the headphones along with the ambient background noise rendered the stimuli presented to the participants inaudible to the experimenter, ensuring that the experimenters were blind to the condition.

### Procedure

#### Experiment 1

Participants faced away from the experimenter and were instructed to raise their hands upon hearing a tone. Tones were initially presented at an SNR of +4 dB (69 dB SPL). The experimenter then iteratively adjusted the level to determine the faintest tone at each frequency that the participant could reliably hear. The experimenter could change the SNR of the signal by steps of .5 dB. The experimenter was blind to the condition (Harmonic/Inharmonic). This procedure is a modified version of the standard clinical audiometry procedure, known as the modified Hughson-Westlake method (Hughson & Westlake, 1944; Carhart & Jerger, 1959). We used this method for efficiency. The experimenter’s blindness to condition precluded differential bias between Harmonic and Inharmonic thresholds.

#### Experiment 2

Singing sessions took approximately 30-40 minutes, and started with a fixed sequence of practice experiments to familiarize participants with the stimuli and instructions. The training was designed to slowly introduce participants to the different elements of the experiment: singing two notes, singing into a microphone, sounds outside the singing range, singing with inharmonic tones, and singing with background noise. Parts of these procedural descriptions are paraphrased or copied from an earlier publication (Jacoby et al, 2019), as the training and setup generally followed the procedure established there.

1. Initial demonstration: The translator provided a Tsimane’ translation of the following sentence: ‘‘The experimenter will make a sound and the translator will copy it. Then, the experimenter will make it again and you will copy it.’’ This verbal description was followed by a demonstration: the experimenter sang two notes at an identical pitch (a unison). The translator then sang back the two notes as best as he could. The experimenter then sang the notes once more, signaling to the participant to sing them back themselves. The process was repeated until participants were comfortable with the flow of the experimental trial format.
2. Training to sing with headphones and microphone: The participant was then familiarized with the use of the headphones and microphone. Most Tsimane’ participants had not used headphones in the past, so the translator helped the participant position the headphones comfortably on their ears, and they were told that the headphones and the microphones were not harmful. The translator provided a Tsimane’ translation of the following sentence: ‘‘Now the computer will make the sound and you will repeat them.’’ The computer played two randomized tones within the singing range for either men or women. The register centers were 320 Hz and 160 Hz for women and men, respectively, and the initial tone for a given participant was randomized to be within +/- 2.5 semitones of that center (chosen from a log uniform distribution). Participants were either presented with the intervals [0 0 -1 -2 -3 -4 0 1 2 3 4] or [0 0 1 2 3 4 0 -1 -2 -3 -4]. The participant replicated the tones by singing. The experimenter waited for the participant to respond before initiating the next stimulus. For this and all subsequent parts of the experiment, tones were 500ms and separated by a 300ms pause, and were pure tones. Pure tones were used for initial training because they are neutral with respect to hypotheses around harmonic vs. inharmonic complex tones.
3. Training to sing continuously: This part of the training session was intended to acquaint the participant with the temporal flow of the experiment. Tones were presented within the participant’s singing range, as in part 2. The conditions (intervals) were presented in one of two ascending/descending orders, counterbalanced across participants ([0, 2, 4, 6, 0, - 2, -4, -6, 0] or [-2, -4, -6, 0, 1, 2, 3, 4, 0]). If a participant had trouble providing timely sung responses, the experimenter and translator repeated the instruction, and this part of the session was repeated.
4. Filter task: participants heard intervals [-4 -3 -2 -1 1 2 3 4] in a random order, in their singing range and the singing range of the opposite sex (order of the ranges was randomized).
5. Training to sing in noise, and training with complex tones: Participants were told that the task would continue in the same way. The experimenter started a 20-minute sample of TE-noise at 65 dB SPL. Participants then completed two blocks of training. In each block, participants heard four trials of either harmonic or inharmonic complex tones in their singing range at 70 dB, followed by four trials at 65 dB SPL, then 60 dB SPL, then 55 dB SPL. The four trials contained +/- 1 and 2 semitone changes, presented in a random order, with random starting notes chosen from the center of the assumed singing ranges (320 Hz for women and 160 Hz for men, starting notes chosen using a log uniform distribution). Participants were not explicitly instructed to listen for tones that were gradually getting quieter, but the gradual nature of this training was intended to show participants that the tones they might hear (and that they were supposed to reproduce) could be quite soft amid the noise. Trials were blocked so that all Harmonic complex tones and all Inharmonic complex tones were presented separately, however experimenters were blind to which block was presented first or second.
6. Main Experiment: The main experiment was separated into four blocks, one for each condition (Harmonic/Inharmonic crossed with Low/High SNR). The experimenter was blind to which block was being presented, and the order of the blocks was randomized. Each block was structured identically except for the type of tone, the SNR, and the f0 range: participants heard 10 intervals, presented in a random order [-4 -3 -2 -1 -.5 .5 1 2 3 4], in either their singing range or the singing range of the opposite sex. Each interval was repeated; the order of repeats was also randomized. This repeat was accidentally omitted for 7 of the 108 Tsimane’ participants due to a coding error. The order of the ranges (male or female singing range) was also randomized. Therefore, each block contained 40 trials total.

Across all parts of the experiment sequence (except for #1 and #2), participants were automatically given four chances to repeat each trial if they did not respond or if it was not possible to extract an accurate pitch measurement from the recording of their response (see below, *Singing Recording and Pitch Extraction,* for possible reasons that a trial might be invalid). After this, the experiment moved on, and the trial was considered a missed trial and a null response for data analysis. In practice, missed trials were infrequent and not evenly distributed across participants. The majority (82%) had none, and this pattern held across both samples: 4/27 US participants and 20/81 Tsimane’ had some missed trials (mean across these participants: 2.4% for US and 6.2% for Tsimane’). We verified post-hoc that group and sub-group differences in pitch and chroma matching persisted (and remained statistically significant) when these participants with missed trials were excluded.

Experimenters were able to monitor the progress of the experiments as the data were collected, by seeing real-time pitch extraction estimates. This enabled the experimenters to determine whether the participants were consistently singing in a way that the computer could not reliably extract pitch (for example, by using excessive vocal fry or by not singing tones long enough).

Trial pacing (stimulus timing and response windows) was fixed and identical across all testing settings.

### Singing Recording and Pitch Extraction

Sung responses were recorded with a Shure SM58 microphone mounted on a microphone stand and connected to a Focusrite Scarlett 2i2 USB sound card. Background noise was minimized at the recording stage as the microphone was positioned approximately 2-4 inches from the singer’s mouth and its cardioid polar pattern and low-sensitivity dynamic transducer attenuates off-axis ambient sounds relative to the close-miked vocal signal. Responses were recorded for 1.43 seconds after the completion of the stimulus (10% longer than the stimulus duration).

Pitch extraction was otherwise identical to that described in Jacoby et al, 2019, and those methods are reproduced in the remainder of this section with minor changes to remove references to experiments in that paper:

We ran an automatic f0 extraction algorithm on the recorded audio immediately following each trial. The f0 extraction algorithm was designed to find long segments of voiced vocal output (corresponding to sung notes) and estimate their fundamental frequency (f0). Trials were considered valid when the algorithm detected exactly two voiced segments (for two note experiments) and the extraction met our quality assurance heuristic criteria (see below). To avoid missing data, if the trial was invalid, we immediately tried to collect another response for the same condition. The trial repeated up to 4 times before being considered a missed trial.

F0 extraction was performed using a custom MATLAB script based on the yin algorithm (de Cheveigne & Kawahara, 2002) with the detection range (the free parameter of the yin algorithm) set to 65.4-523.2 Hz for male participants and 130.8-1046.4 Hz for female participants. Yin outputs the estimated fundamental frequency for each time point within the recording, and an aperiodicity value between 0 and 1 (ratio of aperiodic to total power) that should be low for periodic sound segments. We performed a sliding average of the aperiodicity value over a 10-ms rectangular window. We then extracted continuous voiced segments that could potentially correspond to sung tones by selecting those segments for which the smoothed aperiodicity value was continuously smaller than a threshold of 0.3. This threshold functions as a reliability filter, as frames where periodicity cannot be reliably estimated, including any degraded by residual background noise, are excluded.

We estimated the amplitude envelope of each extracted audio segment as the maximum absolute value of the waveform within a 10 ms rectangular sliding window. This envelope was used for the remaining steps in the note finding procedure. We first located peaks in the envelope using MATLAB’s findpeaks function, and then located the beginning and the end of the voiced segment corresponding to each peak.

To locate the end of the segment corresponding to a peak, we found the earliest point in time t^end^ that satisfied the conditions of (a) having a level of -22 dB or lower relative to the level of the peak and (b) the level of all points within 195 ms after t^end^ (within the window of [t^end^, t^end^ + 195]), remained smaller than the -22 dB threshold. Similarly, we detected the beginning of the segment t^beg^ as the latest time point before the peak where the level within the window ([t^beg^-25, t^beg^]), was lower than -22 dB relative to the level of the peak. To ensure accurate f0 extraction under our field conditions, we further processed the extracted voiced segments using additional heuristics that proved effective in pilot experiments. We found that sung notes were almost always longer than 200 ms, and that shorter voiced segments were usually due to detection errors. We therefore filtered the resulting set of segments by ignoring those with an overall duration of less than 185 ms. We also noticed that participants were less stable in f0 near the beginning and end of a sung note. We therefore discarded the first and last 75 ms of each segment and computed the median of yin’s f0 estimates within the trimmed segment. We also noticed that certain participants sang in a way that produced intermittent octave errors within the produced note, sometimes due to a ‘‘cracked voiced’’ that produced unstable f0 detection, and sometimes due to low SNR. We eliminated such unstable segments by excluding segments where more than 1/3 of the duration of the trimmed segment had an f0 estimate that was more than 6 semitones away from the median. The trial counted as valid when these procedures resulted in two voiced segments, and the median f0s of the trimmed segments were taken as the response. Otherwise, the trial was considered invalid.

### Experiment Locations

Experiments in Tsimane’ were run in four villages: Mara, Moseruna, Majal and Nuevo Mundo. All villages can be reached by a 2.5-4-hour drive from the town of San Borja (the largest town in the region where the Tsimane’ live) if recent weather has been dry. In practice they are often difficult to reach. For example, Mara and Moseruna were not accessible by truck during our fieldwork trip in 2025; all data from Mara and Moseruna were collected in 2024.

Tsimane’ villages do not have enclosed, soundproof spaces, so all experiments were run outside. We took every effort to ensure that experiment locations were quiet and distraction-free. Additionally, ambient noise from the environment was masked by the 65 dB SPL background noise of the main experiments. Previous comparisons between Tsimane’ and US participants have attempted to equate testing conditions by testing US participants in public spaces with ambient background noise (McPherson et al., 2020; McPherson-McNato et al., 2026). Because the experiments in this paper presented stimuli in background noise at 65 dB SPL that as a side effect masked external sources of noise, we deemed it acceptable to run in-person experiments in the US in quiet indoor rooms. Experiments were conducted on the MIT campus in Cambridge, Massachusetts, and the Purdue University campus in West Lafayette, Indiana.

### Study Timeline

In-person experiments were conducted in July-August 2023, 2024, and 2025 for Tsimane’, September 2023-December 2024 for US participants. Online data were collected in March 2026.

### Demographic Survey in Tsimane’

Every Tsimane’ participant completed a demographic survey as part of their experiment session. Using a subset of 16 survey questions related to economic and cultural exposure, we split Tsimane’ participants into two groups. The response to each question was scaled from 0 to 1, and we created an index score for each participant that could be between 0 and 16. All questions are in Supplemental Table 1. The factors we assessed in our survey, including variables such as access and time spent in local towns, access to electricity and technology, contact with local churches and Christian music, were selected from longitudinal surveys and ethnographic data in the Tsimane’ (Leonard et al., 2015; Godoy, 2025), along with our observations of ongoing changes in the region during our work there in the past decade. This approach was inspired by the idea of ‘market integration’, a concept from anthropology referring to lifestyle influences on Indigenous people arising from outside their culture (Lu, 2007).

We used a median split of these scores to divide our 81 Tsimane’ participants into two groups: More Globally Integrated Tsimane’ (N=42, Female=12, mean age = 27.0, S.D.=10.5) and Less Globally Integrated Tsimane’ (N=39, Female=22, mean age=30.2, S.D.=11.2). We refer our two groups of Tsimane’ as ‘More Globalized’ and ‘Less Globalized.

## 3. Statistical Analysis

### Experiment 1

Four detection thresholds were measured per condition, and the three lowest (best) thresholds were averaged for analysis.

### Experiment 2

Direction accuracy was defined as the proportion of sung responses with the same direction (up/down) as the stimulus. It is also possible to analyze the accuracy of the sung response interval. This analysis provides some further support for the primary hypotheses of this paper and is provided in Supplemental Fig. 2.

Chroma and Pitch differences were defined as the difference between a stimulus tone and a response note modulo 12. We analyzed each individual note independently, and split responses into two groups that were analyzed separately: any response that was more than 6 semitones away from the stimulus was analyzed for Chroma matching, and any that was within 6 semitones was analyzed for Pitch matching. Results were qualitatively similar if we simply assigned all responses to tones in the participant’s sex-assigned singing range to the Pitch matching analysis and all those in the range of the opposite sex to the Chroma matching analysis.

We summarized results using histograms over the chroma or pitch difference spanning -6 to 6, and broken into 13 bins: half-semitone bins at the extremes and 11 one-semitone bins in between -5.5 and 5.5. These analysis choices were inherited from a previous publication and were fixed a priori (Jacoby et al., 2019). A difference of 0 corresponds to the response having the same chroma or pitch as the stimulus. Error bars for histograms and null distributions were obtained by bootstrapping. To calculate error bars, we generated 10,000 bootstrap samples, sampling participants with replacement. To calculate null distributions, we used a bootstrap procedure, sampling with replacement 10,000 times, but shuffling the stimulus and response randomly for each participant. To compare across groups, we analyzed the value of the histogram bin around 0 semitones for both pitch and chroma using ANOVAs.

To ensure the robustness of the results to the analysis choices, we recomputed all key comparisons using four alternative outcome definitions: matching-tolerance half-widths of 0.5 (the value used in the main results figures), 1.0, 1.5, and 2.0 semitones. The harmonic advantage in chroma matching and the US-vs-Tsimane’ group differences in chroma and pitch matching were consistent in direction across all definitions, but tended to weaken and lose statistical significance as the bin width increased, consistent with the idea that these effects have a resolution of about a semitone, such that larger bin widths start to mix data from responses that are matched to the stimulus with data from responses that are not matched. The globalization effect (More- vs. Less-Integrated Tsimane’) in pitch matching remained statistically significant across all four bin widths, while the corresponding effect in chroma matching was directionally consistent but did not reach significance under the larger bin widths.

The results were also robust to the 6-semitone cutoff used to separate responses into pitch and chroma matching analyses. Specifically, we reanalyzed results using cutoffs of 4, 5, 6 (the value used for the main results figures), 7, and 8 semitones. The US participants showed significantly greater pitch and chroma matching than Tsimane’ participants at every cutoff (pitch: all t > 2.62, all p ≤ .010; chroma: all t > 2.66, all p ≤ .009). The globalization effect (More vs. Less Integrated Tsimane’) on pooled pitch and chroma matching held at the 6-semitone cutoff used in the main analysis (p=.038) and across nearby cutoffs, but dropped below the .05 significance threshold at the most extreme cutoff tested (p=.069 at 8 semitones).

Together, these analyses indicate that our central conclusions are not specific to the exact bin width or classification cutoff used for the analysis, though they point to a resolution of about a semitone in pitch and chroma matching singing behavior.

Mixed-design ANOVAs were used to test for main effects and interactions between the effects of stimulus condition and participant group. Bayesian analyses were conducted using the bayesFactor MATLAB toolbox (Krekelberg), which implements Bayes-factor hypothesis testing for ANOVA-type designs, including models with fixed and random effects and repeated-measures structures.

As a measure of data reliability, we computed a continuous vocal-reliability index derived from the filtering task. This did not differ between groups and did not account for group differences in pitch or chroma matching, or direction accuracy.

## Data and code availability

Source data is available in this repository: https://osf.io/uwjv3/overview?view_only=e814c93f11024232a26c812e8460b523

## Acknowledgements

The authors thank Tomás Huanca and Esther Conde of the Centro Boliviano de Investigación Socio-integral (CBIDSI) for operational support in Bolivia, Spanish-Tsimane’ translators Elias Hiza Nate, Manuel Roca Moye, and Santos Mosua Semo, drivers Hardy Roca Velarde and Pastor “Gatillo” Roca.

## Author Contributions

Conceptualization: MJM, JHM. Formal Analysis: MJM. Funding Acquisition: MJM, JHM. Investigation: MJM, EAU, AJS, OH. Methodology: MJM, JHM. Project Administration: MJM, EAU, JHM. Resources: MJM, JHM. Software: MJM. Supervision: MJM, JHM. Validation: MJM. Visualization: MJM. Writing – original draft: MJM, JHM. Writing – review & editing: MJM, EAU, AJS, OH, JHM.

## Supplemental Information

### Supplemental Experiment 1

#### Participants

39 participants were enrolled in Supplemental Experiment 1. Recruitment was limited to participants from the US, and any participant with self-reported hearing loss was excluded from the recruitment pool (both criteria were applied using built-in Prolific.com filters). 30 participants (Female=17, mean age=42.6, S.D.=13.9) performed better (i.e., had lower thresholds) than the group mean + one standard deviation, and were included in further analysis.

#### Sample size

We estimated the effect size using a previous experiment that collected a wider range of SNRs and that showed an effect size of d=.86 for a -4.5 dB SNR (-14.5 dB SNR per component) condition. To be adequately powered to detect an effect of this size (at a .05 significance threshold, 95% of the time) we needed 21 participants to power a two-sided Wilcoxon signed-rank test. We collected more data than this target number to enable us to exclude poorly performing participants (using the cutoff described above).

#### Procedure

We measured standard two-tone up/down pitch discrimination with tones presented in noise. On each trial, participants heard two notes, each embedded in masking noise. Participants were asked whether the second note was higher or lower than the first. Adaptive tracks (3-down, 1-up) were initialized at one semitone, and the pitch difference was changed by a factor of 2 for the first four reversals, and a factor of √2 for the final six reversals. Because previous experiments demonstrated that accurate discrimination would be difficult at the lower SNRs for the inharmonic condition, despite tones being audible, we capped possible f0 differences of adaptive tracks at 4 semitones (McPherson et al., 2022). If participants completed three trials incorrectly at this f0 difference, the adaptive track ended early and the threshold was conservatively recorded at 4 semitones (24.99%). Participants completed two adaptive runs per condition, and the order of the tracks was randomized for each participant.

Supplemental Experiment 1 was run online using the platform Prolific (www.prolific.com), we did not have direct control over the sound presentation. However, previous experiments have found that, provided steps are taken to ensure sound quality and participant compliance, there is comparable performance for pitch discrimination in noise with in-lab and in-person data collection (McPherson et al., 2022), and more generally for a range of psychoacoustic tasks (Woods & McDermott, 2018; McWalter & McDermott, 2019; McPherson et al., 2020; McPherson & McDermott, 2020; Traer et al., 2021). We replicated these steps here. First, all participants completed a headphone pre-check to ensure they were wearing headphones or earphones (Woods et al., 2017). Any participant who failed this test did not continue to the main experiment. The experiment also began with a level calibration step during which participants could adjust the sound levels to a comfortable level. For this experiment the absolute level of the tones was less important than the SNRs. Sound was sampled at 44.1 kHz.

#### Analysis

Thresholds were estimated by taking the geometric mean of the f0 differences at the final six reversals of the adaptive track (in semitones). For inharmonic tones, f0 differences were calculated using the generative f0 (i.e. the harmonic f0 that we started with before jittering the component frequencies). The f0 frequency component was never jittered as part of our procedure. Data distributions were non-normal so we used non-parametric tests to perform pairwise comparisons of Harmonic and Inharmonic conditions at the same SNR.

**Supplemental Table 1:** Demographic Survey Questions.

|  | Question | Possible Answers |
| --- | --- | --- |
| 1 | Have you ever lived in any town outside your community for more than six months? | 0=no, 1 = yes |
| 2 | How often do you visit San Borja, Yucumo, Trinidad, or other larger cities? | 0=never, 1=once a year, 2=once a month, 3= every week or more |
| 3 | Do you own a motorcycle, car, or motorized canoe? | 0=never, 0.5=used to, 1=yes |
| 4 | How far is your home community to the towns of San Borja or Yucumo (whichever is closest)? | 0=2 days by canoe, 4+ hours by car, 1=1 day by canoe or 2+ hours by car, 2=Less than 2 hours by car, 3=less than an hour by car |
| 5 | Have you worked for a company in the area in the last year (cattle, logging, trader)? | 0=never, 1=occasionally, 2=often |
| 6 | How often do you buy, sell, or trade goods? | 0=never, 1=once a year, 2=once a month, 3= every week or more |
| 7 | Do you speak Spanish? | 0=no; 0.5=little; 1=fluent |
| 8 | What is the highest grade of schooling you have completed? | 0-12 for none to 12<sup>th</sup> grade |
| 9 | Do you have electricity in your home (*i.e., is your home connected to the electrical grid*), a solar panel, or a generator? | 0=never, 0.5=used to, 1=yes |
| 10 | Do you have a working television in your home? | 0=never, 0.5=used to, 1=yes |
| 11 | Do you own a working cell phone? What type? | 0=never, 1= used to, 2=yes, not a smartphone, 3=yes, a smartphone |
| 12 | Do you have a working radio in your home? | 0=never, 0.5=used to, 1=yes |
| 13 | How often do you attend church? | 0=never, 1=once a year, 2=sometimes, 3=every Sunday or more |
| 14 | How many Christian hymns do you know how to sing (either in Tsimane' or Spanish)? | 0=None, 0.5=Some, 1=Many |
| 15 | Do you sing? If yes, where do you sing? (at home, at church, with friends) If yes, how often do you sing?<br>*Note: for questions 15 and 16, we found that asking participants where they sang or played instruments helped to elicit accurate estimates of frequency.* | 0=never, 1=once per year, 2=between 1-3 times per month, 3=once a week, 4=more than once a week |
| 16 | Do you play a musical instrument? If yes, where do you play? (At home, at church, with friends). If yes, how often? | 0=never, 1=once per year, 2=between 1-3 times per month, 3=once a week, 4=more than once a week |

**Supplemental Figure 1:**
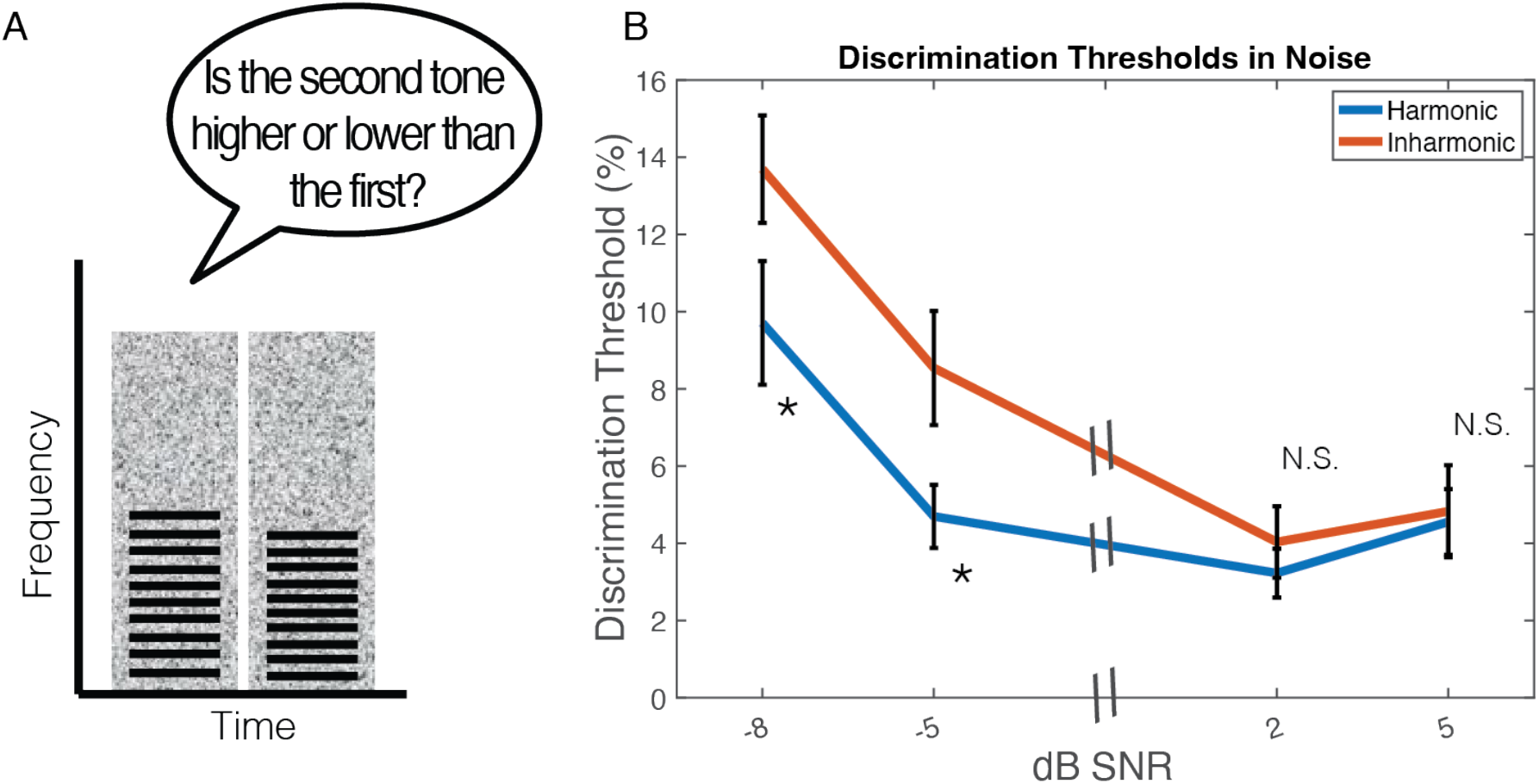
Results of Supplemental Experiment 1. Previous experiments exploring discrimination of harmonic and inharmonic tones in noise suggested that at low SNRs, we would observe a harmonic advantage, but that this advantage would not be present at high SNRs (Carlyon & Stubbs, 1989; McPherson et al., 2022). The aim of this Experiment was to validate the choice of SNRs for Experiment 2, where we planned to test individuals at two SNRs, one yielding better thresholds for harmonic than inharmonic tones in US listeners, and the other yielding comparable thresholds. A. Schematic of trial structure. In each trial, US participants heard two noise bursts, each of which contained a tone, and were asked whether the second note was higher or lower than the first tone. Tones were either both Harmonic or both Inharmonic. F0 differences were adaptively changed to find the discrimination threshold. Participants completed two adaptive tracks per condition for Harmonic vs. Inharmonic conditions at 4 SNRs: -8, -5, 2, and 5 dB SNR. B. Results. We observed significantly lower thresholds for Harmonic than Inharmonic tones at -8 dB SNR (*p*=.031,*z*=-2.16) and -5 dB SNR (*p*=.039, *z=*-2.07), but no significant difference in thresholds at higher SNRs (2 dB SNR, *p*=.98, *z*=-0.03,and 5 dB SNR, *p*=.57, *z*=0.56. For inharmonic tones, f0 differences were calculated using the generative f0 (i.e. the harmonic f0 that we started with before jittering the component frequencies). The f0 frequency component was never jittered as part of our procedure. Error bars denote standard error of the mean. Asterisks denote statistical significance of a Wilcoxon signed-rank test between Harmonic and Inharmonic conditions, *=p<.05. These results support the choice of -8 and 5 dB SNR for Experiment 2.

**Supplemental Figure 2:**
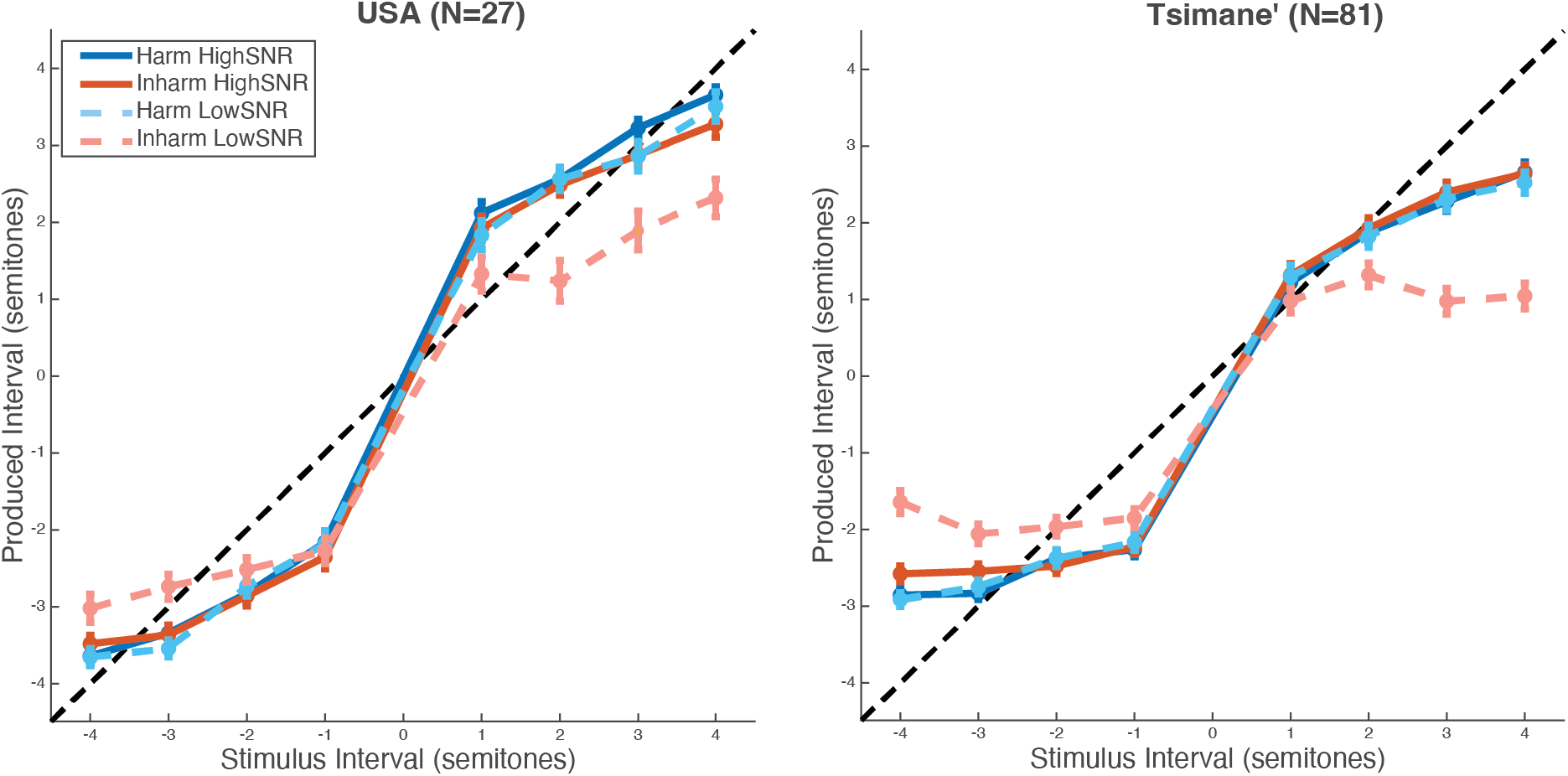
Sung Interval Accuracy. Results of an analysis examining the relationship between stimulus and response intervals. Each data point represents the mean reproduced interval for a particular stimulus interval. Error bars are standard error of the mean.

**Supplemental Figure 3:**
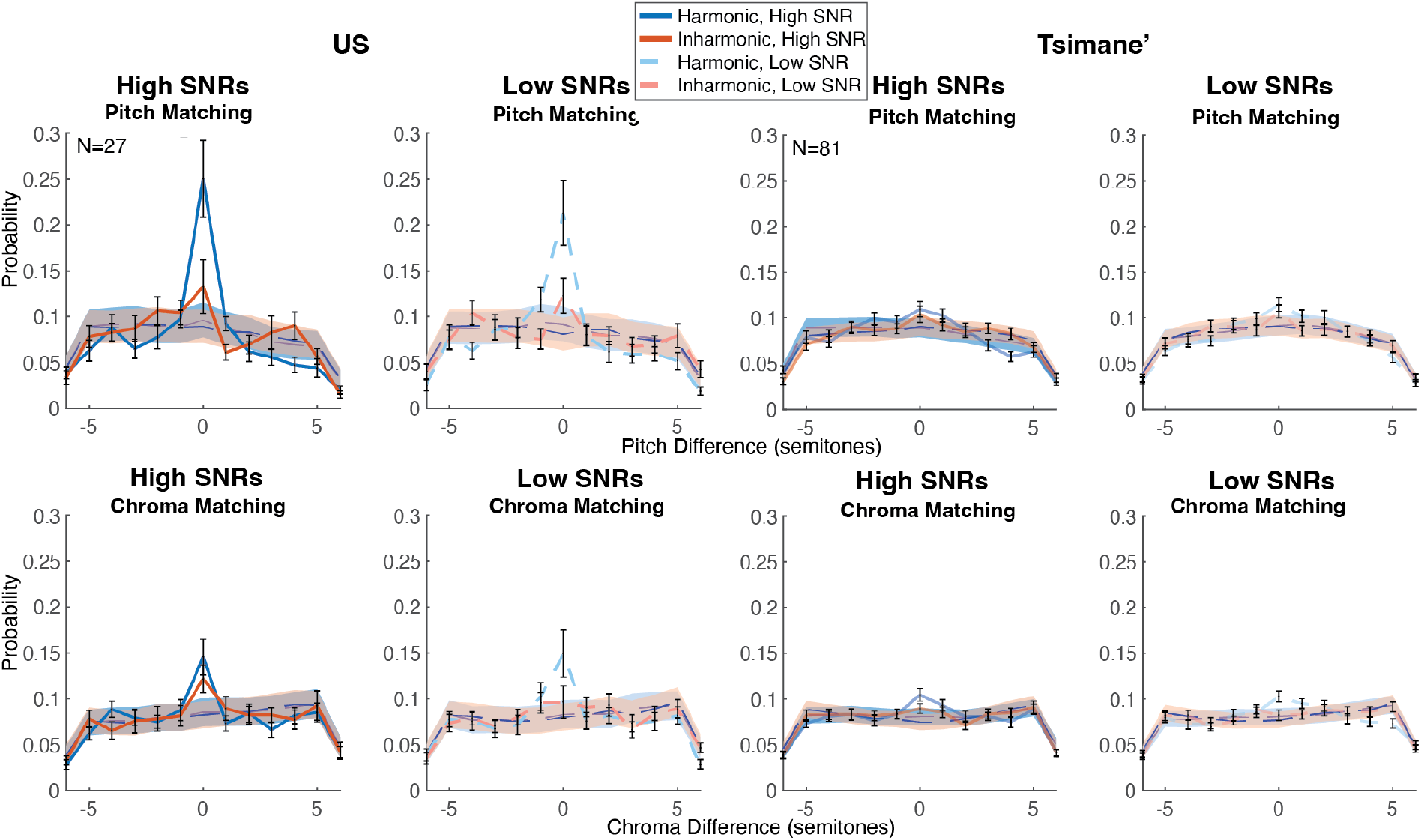
Pitch and Chroma Matching Results for High and Low SNRs. Histograms of stimulus-response pitch (top row) and chroma (bottom row) differences High and Low SNR conditions, separated by participant group. Error bars plot SEM across participants, calculated via bootstrap. The shaded regions denote the null distributions, computed from histogram of permuted datasets where stimulus-response correspondence was randomized.

**Supplemental Figure 4:**
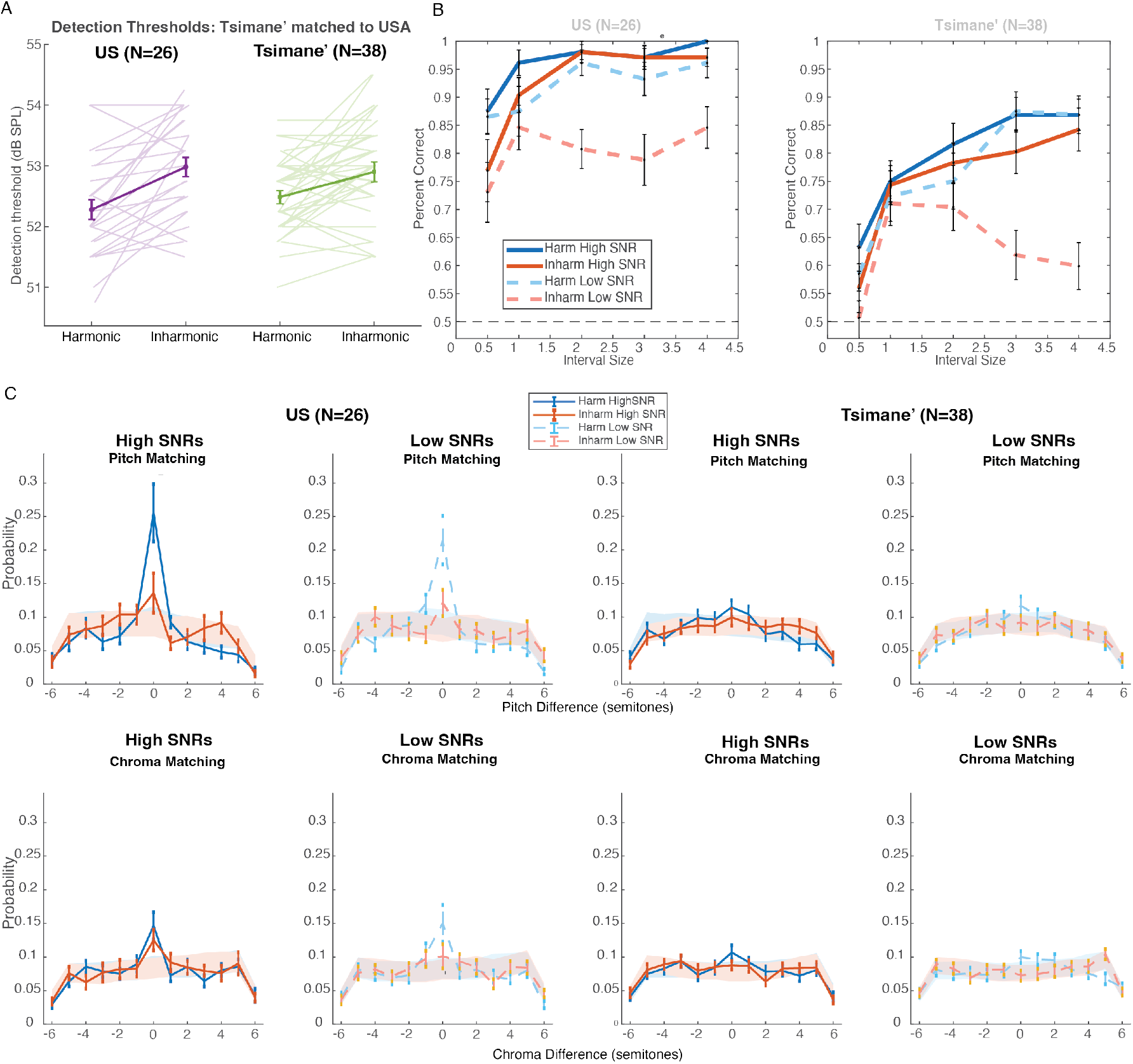
Threshold-matched subsample reproduces the main cross-cultural effects. To test whether group differences in detection thresholds (Fig. 1e) could account for the effects observed in Experiment 2, we split the Experiment 1 data in half for each participant, used one half to select a subset of Tsimane’ participants with detection thresholds closely matched to the US group, and used the second half to confirm the match (plotted). (A) Individual (thin lines) and group-average (thick lines with error bars) detection thresholds for Harmonic and Inharmonic tones in the threshold-matched US (N=26) and Tsimane’ (N=38) subsamples; average thresholds did not differ significantly between groups (t(62)=0.15, p=.88, BF10=0.26). (B) Direction accuracy (percent correct) as a function of interval size for the matched US and Tsimane’ subsamples, plotted separately for Harmonic and Inharmonic tones at High and Low SNR. As in the full sample, both groups showed comparable performance for Harmonic and Inharmonic tones at High SNR but a pronounced advantage for Harmonic over Inharmonic tones at Low SNR, with no Group interactions (all p>.50). (C) Histograms of stimulus-response pitch differences (top) and chroma differences (bottom) for High and Low SNR conditions in the matched subsamples. Shaded regions denote null distributions from permuted (stimulus-response shuffled) data; error bars show bootstrapped SEM across participants. As in the full sample, US participants showed clear peaks at 0 semitones indicating pitch and chroma matching, while matching was comparatively weak in the Tsimane’ subsample. Group differences in direction accuracy (F(1,62)=21.92, p<.0001, ηp²=.26), chroma matching (F(1,62)=7.02, p=.010, ηp²=.10), and pitch matching (F(1,62)=9.41, p=.003, ηp²=.13) all remained significant after matching for detection sensitivity, indicating that the cross-cultural differences reported in the main text are not attributable to differences in overall auditory thresholds between groups.

**Supplemental Figure 5:**
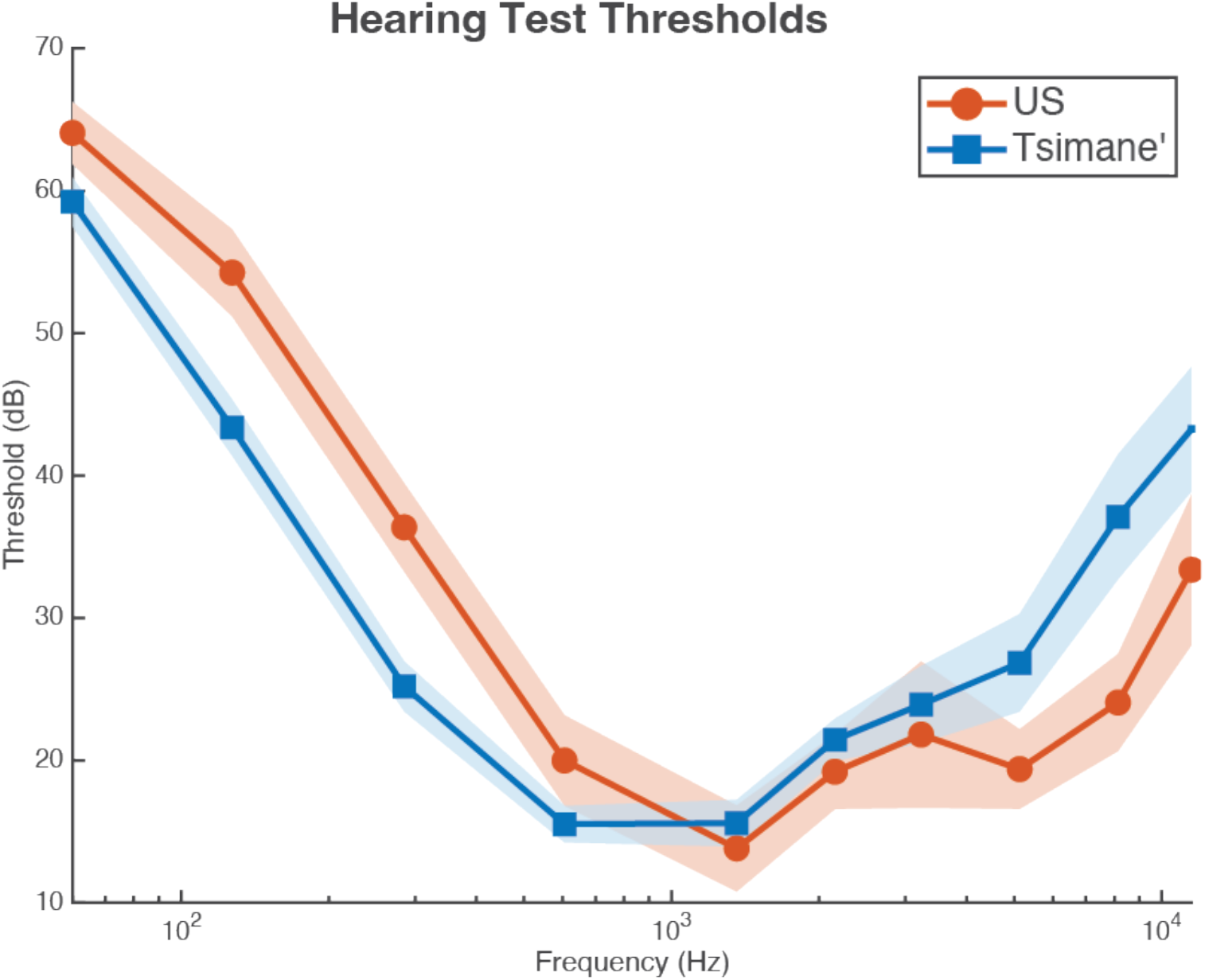
Hearing test thresholds. Results of hearing screenings in US and Tsimane’ participants. Participants were seated facing away from the experimenter and asked to raise their hand every time they heard a tone. Participants heard pure tones (presented diotically) at 60, 127, 285, 605, 1358, 2155, 3229, 5126, 8137, 11500 Hz. The tones were presented in 10 randomly ordered blocks, and the experimenter adjusted the intensity of the tones to determine the faintest tone the participants could hear at each frequency. The differences in thresholds between groups are consistent with prior work (Jacoby et al., 2019) and plausibly reflect differences in the average spectrum of the ambient background noise between testing sites, which despite being attenuated by the closed headphones inevitably has some effect on detection thresholds. Shaded areas show standard error of the mean.

**Supplemental Figure 6:**
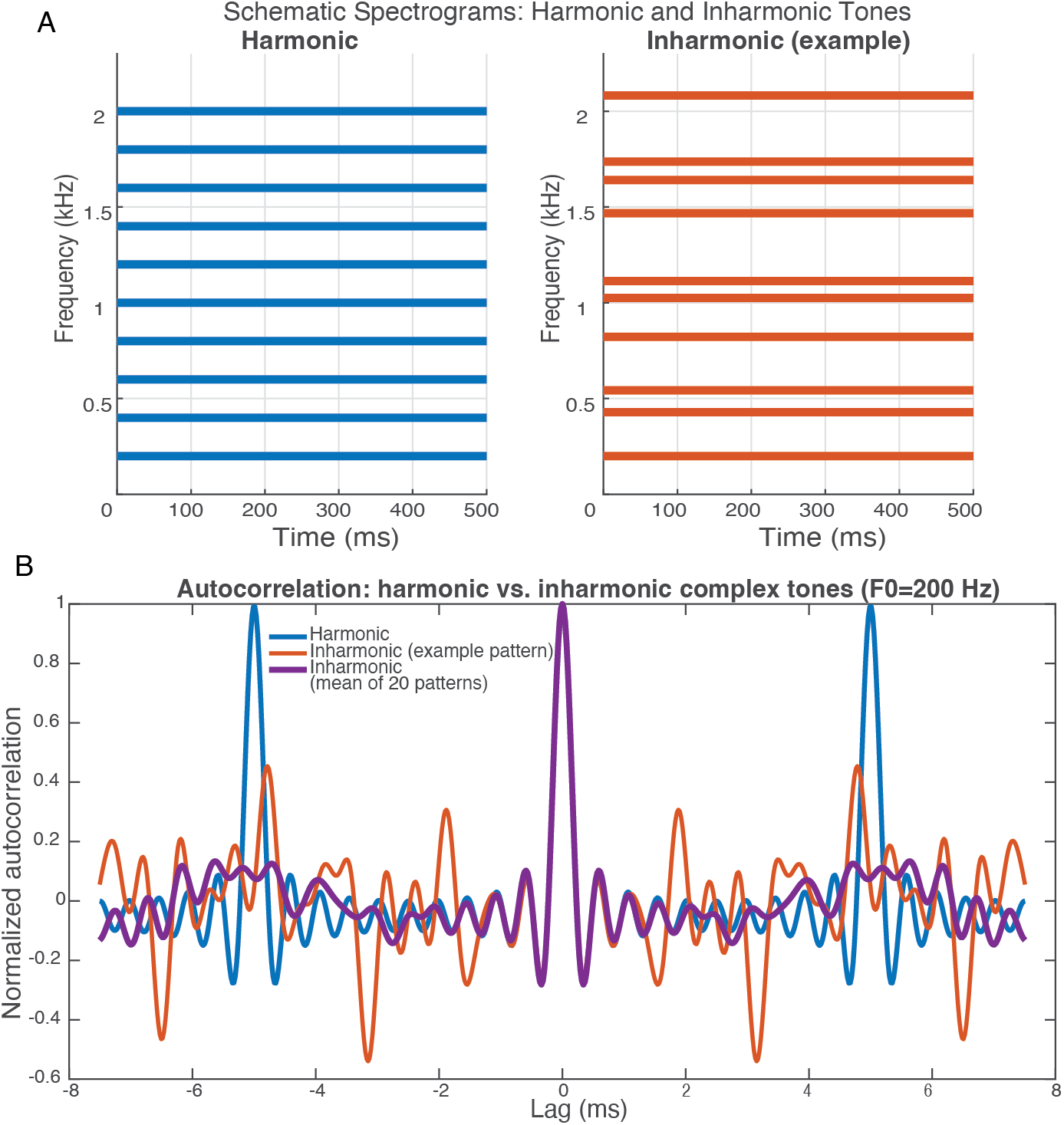
Stimuli visualizations. (A) Schematic spectrogram of a Harmonic and Inharmonic tone, each with 10 equal amplitude harmonics. (B) Autocorrelation function of a harmonic tone, an example inharmonic tone, and the mean autocorrelation of all 20 inharmonic patterns used throughout the experiment. The 20s were selected from an initial set of 100,000 possible jitter patterns because they minimized the peak of the autocorrelation.

